# Hearing loss in juvenile rats leads to excessive play fighting and hyperactivity, mild cognitive deficits and altered neuronal activity in the prefrontal cortex

**DOI:** 10.1101/2023.08.23.554422

**Authors:** Jonas Jelinek, Marie Johne, Mesbah Alam, Joachim K. Krauss, Andrej Kral, Kerstin Schwabe

## Abstract

**Background:** In children, hearing loss has been associated with hyperactivity, disturbed social interaction, and risk of cognitive disturbances. Mechanistic explanations of these relations sometimes involve language. To investigate the effect of hearing loss on behavioral deficits in the absence of language, we tested the impact of hearing loss in juvenile rats on motor, social, and cognitive behavior and on physiology of prefrontal cortex.

**Methods:** Hearing loss was induced in juvenile (postnatal day 14) male Sprague-Dawley rats by intracochlear injection of neomycin under general anaesthesia. Sham-operated and non-operated hearing rats served as controls. One week after surgery auditory brainstem response (ABR) measurements verified hearing loss or intact hearing in sham-operated and non-operated controls. All rats were then tested for locomotor activity (open field), coordination (Rotarod), and for social interaction during development in weeks 1, 2, 4, 8, 16, and 24 after surgery. From week 8 on, rats were trained and tested for spatial learning and memory (4-arm baited 8-arm radial maze test). In a final setting, neuronal activity was recorded in the medial prefrontal cortex (mPFC).

**Results:** In the open field deafened rats moved faster and covered more distance than sham-operated and non-operated controls from week 8 on (both p<0.05). Deafened rats showed significantly more play fighting during development (p<0.05), whereas other aspects of social interaction, such as following, were not affected. Learning of the radial maze test was not impaired in deafened rats (p>0.05), but rats used less next-arm entries than other groups indicating impaired concept learning (p<0.05). In the mPFC neuronal firing rate was reduced and enhanced irregular firing was observed. Moreover, oscillatory activity was altered, both within the mPFC and in coherence of mPFC with the somatosensory cortex (p<0.05).

**Conclusions:** Hearing loss in juvenile rats leads to hyperactive behavior and pronounced play-fighting during development, suggesting a causal relationship between hearing loss and cognitive development. Altered neuronal activities in the mPFC after hearing loss support such effects on neuronal networks outside the central auditory system. This animal model provides evidence of developmental consequences of juvenile hearing loss on prefrontal cortex in absence of language as potential confounding factor.

## 1. Introduction

Hearing loss is the world’s leading disability, affecting 450 million people worldwide, of which 34 million are children (OMS O, 2021). In the elderly, the current focus is on the association with dementia (Livingston et al., 2020; Nadhimi & Llano, 2021), whereas hearing loss in early childhood is discussed in relation to communication and educational deficits (AuBuchon et al., 2015; Idstad & Engdahl, 2019; Tager-Flusberg, 2015; Teasdale & Sorensen, 2007; Tomblin et al., 2015), as well as hyperactivity and socio-emotional problems (Altshuler, 1978; Bigler et al., 2019; Lieu et al., 2020). The link between hearing and higher cognitive functions is also supported by the fact that not only spoken language outcomes but also cognitive abilities improve after hearing restoration with cochlear implants (AuBuchon et al., 2015; Lazard et al., 2012; Wong et al., 2017). However, deficits in complex cognitive skills persist in some children despite early intervention for hearing impairment(Kronenberger et al., 2013; Marschark et al., 2007). A key role in language in the link between hearing loss and risk of abnormal cognitive functions has been hypothetized (Hall et al., 2017). If true, this implies that hearing loss in absence of language, as observed in animals, should not be related to cognitive deficits.

A rodent model provides the opportunity to investigate the causality between hearing loss and cognitive function in juvenile rats and their development in a controlled environment, without the influence of language. Longitudinal studies of adult rat behavior after hearing loss showed only a mild effect on learning the concept of a radial arm maze task, transient hyperactivity, and no effect on social interaction (Johne et al., 2022). Early life insults, however, not only induce compensatory changes but also lead to compromised functional integrity of neural networks developing after birth with behavioral, neurophysiological and neuroanatomical consequences (Harich et al., 2008; Helgers et al., 2020), with the influence of hearing playing an important role (Kral et al., 2005; Kral & Sharma, 2012). The strong impact of experience on neuronal circuits leads to developmental critical periods for therapy of hearing loss (Kral et al., 2019). Hearing loss in juvenile age therefore has more severe or different consequences than hearing loss in adulthood and affects also outcomes of hearing restoration in later life. Social interaction also changes with age. While juvenile rats engage in rough and tumble social play behavior, peaking at 30-40 days of age, this behavior declines in adult rats (Papilloud et al., 2018; Thor & Holloway, 1984). The prefrontal cortex (PFC) is of particular interest in this regard, as it is critical for higher cognitive and executive functions, as well as for normal socio-emotional functioning (Teffer & Semendeferi, 2012). Furthermore, it is anatomically and reciprocally connected to virtually all sensory and motor systems, as well as to a variety of subcortical structures (Fassbender & Schweitzer, 2006; Li et al., 2015; Miller & Cohen, 2001; Russo & Nestler, 2013). In rats with induced hearing loss in adulthood, electrophysiological recordings in the medial PFC (mPFC) showed substantially altered neuronal activity, both at the neuronal and network level (Johne et al., 2022).

In this study, hearing loss was induced in rats at postnatal day (PND) 14 by intracochlear injection of neomycin, followed by longitudinal behavioral testing of motor activity and social interaction, as well as spatial learning and memory. These rats were in adult age (i.e. 6 months after deafening) subjected to final electrophysiological recordings of single unit (SU) activity and local field potentials (LFP) in the medial PFC (mPFC) and coherence with the sensorimotor cortex (SMCtx). With respect to SU, we distinguished between the two main types of neurons in the mPFC: excitatory pyramidal cells, which make up ∼80% of all neurons and project across brain areas, and interneurons, which are mainly inhibitory and whose axons remain within a circumscribed cortical area (Diester & Nieder, 2008).

## 2. Material and Methods

### 2.1 Animals

Adult male Sprague-Dawley rats (Charles River Laboratories; n=29) were housed in Makrolon Type IV open cages in groups of two or three animals in the same experimental subgroup under controlled environmental conditions (22±2°C; 55±10% humidity; 10/14h dark/light cycle with lights on at 06:00 am). A standard rodent diet (Altromin, Lage, Germany) was fed daily at 14-16 g per animal from 200 g body weight on to maintain uniform weight gain without fattening. Tap water from rodent drinking bottles was available at all times. To ensure the welfare of the animals, they were weighed and clinically examined at least twice a week and visually inspected daily. The animals’ health monitoring was performed according to FELASA recommendations using sentinel screening.

All experiments conducted were carried out strictly in accordance with the guidelines of the EU directive 2010/63 and were approved by the local animal ethics committee, including power analysis and blinded and randomized study design (Lower Saxony State Office for Consumer Protection and Food Safety, AZ 18/2874). Every effort was made to minimize animal numbers and ensure the best possible animal welfare according to the 3R-principle.

### 2.2 Study Design

After birth, rats were housed with their mothers in groups of eight pups per litter until weaning at postnatal day 21 (PND 21). At PND 14, litters were randomly divided into three groups: (1) the deafened group (n=13) in which hearing loss was induced by intracochlear application of neomycin, (2) the sham-deafened group (n=8) with the same surgical procedure without application of neomycin, and (3) the non-operated control group (n=8). After weaning at PND 21 (i.e., one week after surgery) the auditory brainstem response (ABR) was measured to verify deafness or normal hearing in the sham-deafened and naïve control rats.

At weeks 1, 2, 4, 8, 16, and 24 after surgery rats were subjected to behavioral tests of locomotor activity (open field), social interaction within pairs of two rats, and balance (Rotarod). In the eighth postoperative week, anxiety behavior was tested once in the plus maze. Training then began in the 4-arm baited 8-arm maze. After reaching a certain performance criterion, maze behavior was tested again at weeks 16 and 24. Social preference was tested at weeks 8, 16, and 24 (Figure 1A).

**Figure 1:**
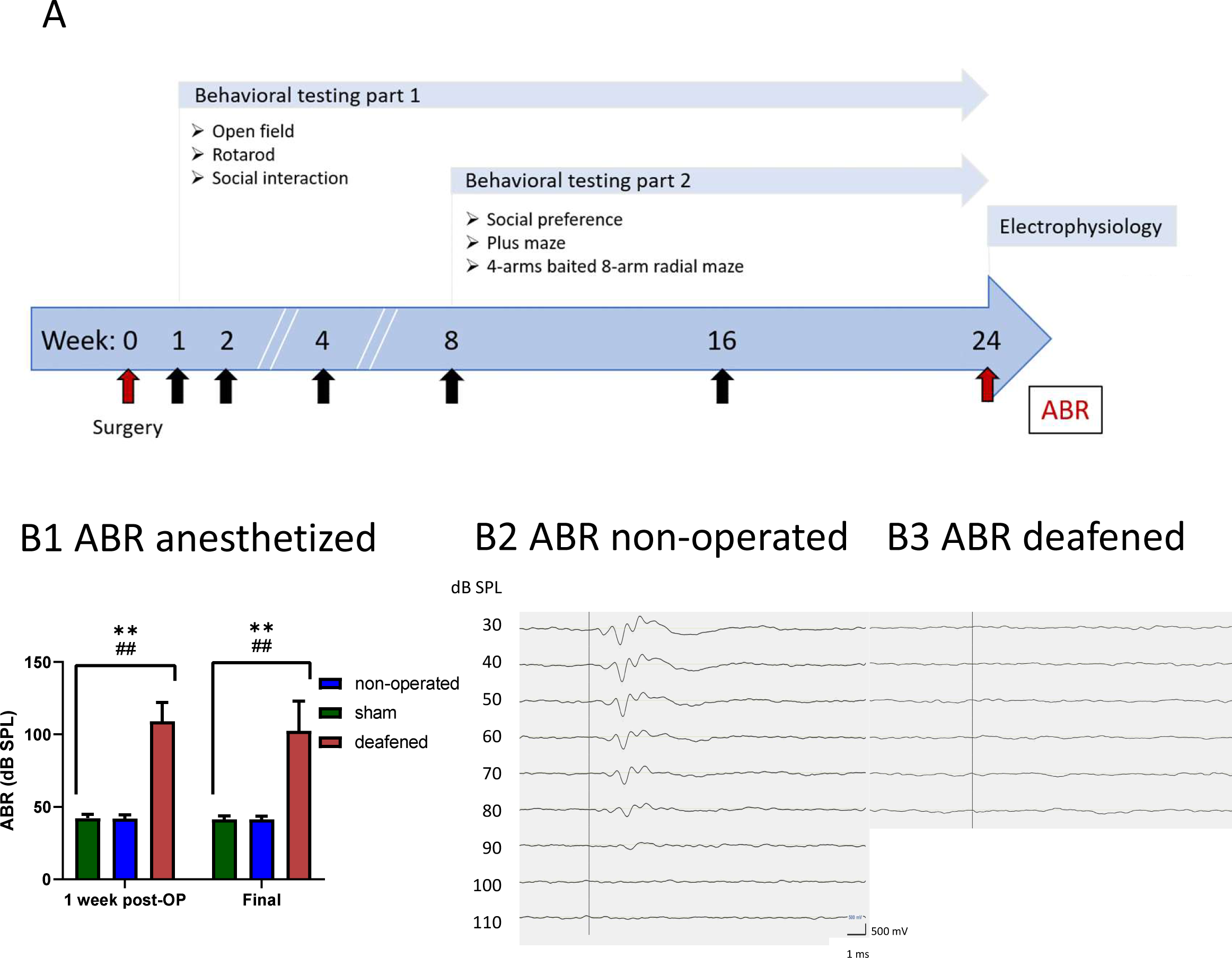
Experimental design and ABR measures. Chronology of the experimental setup **(A)**. Effects of the deafening surgery to the auditory brainstem response of non-operated, sham, and deafened rats **(B1)** with data shown as means ± SEM post OP and at final measurements 24 weeks after surgery. Differences between deafened rats and non-operated controls are shown as asterisks (*) and between deafened and sham as hashtag (#) (*p < 0.05, **p < 0.01 after ANOVA). In addition, archetypal examples of ABRs from a non-operated rat **(B2)** and a deafened rat **(B3)** are shown.

To assess possible age-related changes ABR measurements were repeated at week 24 just before final electrophysiological recordings bilaterally in the mPFC. The exact location of the electrophysiological recordings in the mPFC was confirmed after histological processing of the brains using Nissl-stained brain sections.

### 2.3 Surgery

Prior to surgery, the entire litter was removed from the mother for the duration of surgery. All rats were weighed and randomly assigned to the deafened, sham-deafened or non-operated group. The surgical procedure was performed under general anesthesia with isoflurane (Isofluran-Piramal, Voorschoten, Netherlands). In addition, the tissue in the postauricular region was infiltrated with local anesthesia (Lidocain-“Welk”, Combustin, Haitlingen, Germany) and systemic pain therapy (5 mg/kg carprofen s.c.) was administered. Surgery followed the procedure described by Jakob et al., (2016). After positioning the animal, a postauricular incision of about 2cm length was made at the level of the external acoustic canal. After blunt preparation to expose the tympanic bulla and the facial nerve, the tympanic bulla was opened with pointed tweezers. The round window membrane was punctured with a Hamilton syringe and 3 x 15 μl neomycin (5 mg dissolved in 1ml water, adjusted to pH 7.4) was injected to induce sensorineural hearing loss. Thereafter, the skin was closed with a skin suture. After performing the same procedure on the opposite side, the animal was placed on a warming mat and examined closely until it was fully awake, at which point all the animals in one litter were reunited with their mother. On the following two days, the rats received systemic pain therapy (2.5 mg/kg carprofen s.c.) and were weighed and clinically evaluated daily for one week. Rats in the sham-deafened group underwent the same surgical procedure, but without opening of the tympanic bulla and without neomycin injection. Non-operated control animals received no anesthesia, surgery, or medication.

### 2.4 Hearing loss assessment

For the hearing assessment the same setup was used as in a previous study (Johne et al., 2022). For ABR measurements three subdermal monopolar needle electrodes were placed (1) medially on the skull at the lateral corner of the eye, (2) behind the ear, and (3) as a grounding electrode in the skin fold on the back. Auditory brainstem responses were obtained from vertex vs. the postauricular electrode by 40 dB preamplification (AMP55, Otoconsult Comp., Frankfurt, Germany) and 60 dB amplification (PNS1, Otoconsult Comp., Frankfurt, Germany). To avoid interfering low frequency signals a second order bandpass filter (100-5000Hz) was applied. Stimuli were generated by Audiology Lab stimulus generation and acquisition system (Otoconsult Comp., Frankfurt, Germany) running on a PC and connected to an AD converter (National Instruments NI-USB-6251, Austin, TX, ZSA) and an analogue USB-controlled attenuator (PNS1, Otoconsult Comp., Frankfurt, Germany) connected to a calibrated DT48 speaker (Beyerdynamic, Heilbronn, Germany) positioned close to the ear. Condensation clicks of 50 µs duration were presented at levels from 10 to 100 dB SPL p.e. in 10 dB steps. The interstimulus interval was 113 ms and clicks were repeated 200 times for each stimulus intensity. Recorded signals were bandpass filtered (100 Hz – 10 kHz). For analysis, 200 sweeps per volume were averaged using custom codes in MATLAB (The MathWorks, Natick, USA). Signals were additionally filtered (300–3000 Hz). ABRs were determined manually by appearance of ABR components 2 – 3 ms after stimulus onset (Johne et al., 2022; Schopf et al., 2014; Tillein et al., 2012).

### 2.5 Behavioral testing

#### Open field

To test locomotion behavior rats were placed in an open field arena (62x62x30 cm) for ten minutes and filmed from above. Locomotor distance and velocity were evaluated automatically (TopScan; TopView Analyzing System 2.0; Clever Sys Inc.).

#### Social interaction

To assess social interaction, two rats of the same experimental group were placed in the open field arena for ten minutes and recorded from above with a fixed camera. The social behavior (play-fighting, following, and total interaction) was analyzed offline using ODLog™ (Macropod Software™ version 2.7.2).

#### Social preference

To study social preference, a rat was placed in the open field for 5 minutes to acclimatize. Thereafter, an empty box was added for 4 minutes, followed by another box containing an unfamiliar rat for a further 4 minutes. The number of interactions with the object alone and the object with the social stimulus partner was scored.

#### Rotarod

Motor coordination was assessed by placing the animals on a rotating rod (10.5x43x43 cm; Rota-rod, Series 8, IITC Life Science, Cambridge, UK). The rod rotated with increasing speed for 60 seconds (initial speed: five revolutions per minute; maximum speed: 15 revolutions per minute), followed by another 60 seconds at a constant speed (15 revolutions per minute). Rats were tested in three consecutive trials, and the time was measured until the rat fell off or the time ended. The best time of all three trials was used for analysis.

#### Plus maze

Anxiety was tested on an elevated plus maze consisting of four elevated arms (12 cm wide, 76 cm long, 80 cm high) connected in a cross-like disposition by a central platform (12 cm x 12 cm). Two opposite arms were closed by walls (27 cm in height), the other two arms were open. For testing, rats were placed on the central platform facing an open arm. During the ten minutes test period the number of entries in open and closed arms, and on the central platform were assessed online via a screen outside of the experimental room.

#### 4-arm baited 8-arm maze

The maze consisted of eight arms (12 x 76 cm) projecting radially from a central platform (35 cm in diameter) with adjacent arms separated by 45° and 80 cm above the floor. A metal cup for reward pellets (dust-free precision pellets, 45 mg purified rodent food, Bio-Serv, Flemington, NJ, USA) was placed at the end of each arm. The maze was set in an experimental room with several external visual cues. The experimenter monitored the movements of the rats via a video camera mounted above the maze and a TV-screen outside the experimental room. The experiments began with a habituation period of two days in which each animal was allowed to freely explore the entire maze. For the test, four randomly selected arms (no more than two adjacent arms) were provided with two reward pellets in each cup. The test began by placing the animal in a random direction in the center of the maze and ended when the animal collected all rewards or after 15 minutes.

The sequence of arms visited was followed from outside the room by video recordings. For analysis, the number of entries to unrewarded arms (reference memory errors (RME)) and the number of re-visited arms during the same trial (working memory errors (WME)) were assessed. Additionally, the order of arm choices was monitored. The training was terminated upon a completion criterion of fewer than three WMEs in three consecutive blocks (each consisting of three runs) was met along with fewer than two RMEs per block.

After the initial training for the completion criterion, rats were retested in the radial maze every 8 weeks. In weeks 16 and 24, animals were tested on two consecutive days with two blocks each (12 trials in total). As a final test, all animals were subjected to a reversed training phase of 4 blocks on two days (i.e., 12 trials) in which rewards were placed in the previously unrewarded arms to investigate the ability to adapt to the new situation. For analysis, the number of RME, WME and the percentage of entries into arms next to the previously visited arm, i.e., 45° turns (number of all turns/number of 45° turns x 100), were analyzed in blocks of three trials.

### 2.6 Electrophysiology

Electrophysiological recordings were performed with a similar setup as previously used in adult deafened rats (Johne et al., 2022). For extracellular measurements of SU neuronal activity and recording of LFPs, rats were anesthetized with urethane (1.4 g/kg in 0.9% NaCl, i.p. ethyl carbamate, Sigma); the depth of anesthesia was regularly checked by pinching the feet and additional doses were administered as needed. The incision site was infiltrated with a local anesthetic, Lidocain (Lidocain-“Welk”, Combustin, Haitlingen, Germany). Rats were placed in a stereotaxic frame, and body temperature was maintained at 37 ± 0.5°C with a warming mat (FHC, Bowdoinham, ME, USA) and monitored permanently by rectal temperature measurement. Small craniotomies were performed at the target coordinates for the mPFC on both hemispheres. The animal was then placed in a Faraday cage to minimize electrical interference. A single microelectrode for extracellular leads (quartz-coated electrode with a platinum-tungsten alloy core (95-5%), a diameter of 80 µm, and impedance of 1-2 MΩ was connected to the Mini Matrix 2-channel version drives headstage (Thomas Recording GmbH, Giessen, Germany). The microelectrode signal was passed through a head stage and then split into SU and LFP components separately. The stainless steel guide needle contacting the cortical surface served as a reference for the microelectrode, as previously reported (Alam et al., 2012; Elle et al., 2020). For SU recording, the signals were bandpass filtered between 500 and 5000 Hz, amplified from x 9500 to 19,000, and sampled at 25 kHz. Additionally, ground wires were clamped at the neck as grounding. Recordings in the mPFC were made at the following coordinates in mm relative to the bregma and ventral to the cortical surface; anterior-posterior (AP) +3.2 and +2.7; mediolateral (ML) ±0.5 and ±0.8; ventral (V) 3.2 to 4.5. Primary SMCtx electrocorticograms (ECoGs) were recorded via a 1-mm-diameter jeweler’s screw positioned on the dura mater above the primary SMCtx region (AP: -0.4 mm; ML: ±2.5 mm). The signal was bandpass filtered (0.5-100 Hz) with a sampling rate of 1 kHz. Data were acquired using the CED 1401 A/D interface (Cambridge Electronic Design, Cambridge, UK). After completion of the experiment, electrical lesions were made at the recording sites to allow histological verification of localization (10 µA for 10 seconds; both negative and positive polarity), as previously described (Jin et al., 2016).

#### SU analysis

Action potentials originating from a single neuron were discriminated using the template matching function of spike sorting software (Spike2; Cambridge Electronic Design, Cambridge, UK). Only well-isolated SUs were included in the analysis, as determined by homogeneity of spike waveforms, separation of spike waveform projections to principal components during spike sorting, and clear refractory periods in inter-spike interval (ISI) histograms. All analyses were performed using custom-written MATLAB functions (Mathworks, Natick, MA, USA) unless otherwise noted.

FR was analyzed by taking the reciprocal of the mean ISI for the entire 100 s of the recording. The dispersion index (DI) of FR, a measure of the variance in firing rate, was calculated by dividing the square of the standard deviation of ISI by the mean ISI (std of ISI2/mean ISI) to compare the variation in their median values. A higher random firing pattern would be expected to increase the diversity of ISI lengths and have a higher DI of FR. A lower DI would indicate less diversity of ISI lengths and more regular activity.

Previous work with extracellular recording studies used waveform durations to classify neurons in the PFC and in other cortical regions into putative pyramidal neurons and putative interneurons (Diester & Nieder, 2008; Elle et al., 2020; Mitchell & Maslin, 2007). These studies revealed that action potentials of pyramidal neurons have longer durations than those of interneurons. The SU activities were therefore classified as either narrow spike (NS) for inhibitory neurons and broad spikes (BS) for glutamatergic as excitatory neurons on spike shape as previously described (Elle et al., 2020). The time between the peak and trough of the spike waveform was measured and the short duration group of neurons (peak to trough time of 0.12-0.36 ms) was considered NS, the long duration neurons (time window >0.36 ms) were considered BS. Waveform duration was determined by measuring the time from the trough (negative deflection) to the peak (positive deflection). All classifications of neurons as NS or BS were based on these values.

**LFP and coherence analysis:** representative epochs of 100 s were used for signal processing in the frequency domain for the LFPs in the mPFC, and SMCtx-EcoGs. A 50 Hz notch filter with finite impulse response and a 100 Hz low-pass filter were used. The spectral power of the mPFC, and SMCtx-EcoGs was derived by discrete Fourier transform with blocks of 1024 samples using a Welch periodogram in a custom Matlab script (MathWorks, Inc.), resulting in a frequency resolution of 0.9766 Hz. The Hanning window function (also called Hann window) was applied to avoid the spectral leakage phenomenon. The relative power indices for each band were calculated using the absolute power in each frequency band and expressed as a percentage of the absolute power. To compare the powers in different frequency bands, the area under the calculated power density spectrum in specific frequency ranges was calculated and averaged for theta (4-8 Hz), alpha (8-12 Hz), beta (12-30 Hz), and gamma (30-100 Hz) frequency bands.

Furthermore, the functional relationships between the SMCtx-EcoGs and the LFPs of the mPFC were estimated by coherence as described by (Halliday et al., 1995). Coherence analyses are used to determine the strength of oscillatory synchronizations in brain networks in various neurological and neuropsychiatric disorders. The coherence of oscillatory signals is a measure of the linear phase and amplitude relationships between signals in the frequency domain. It is a finite measure with values from 0 to 1, where 0 indicates no linear relationship and 1 indicates a perfect linear relationship.

### 2.7 Histology

Immediately after electrophysiological recording rats were injected with a lethal overdose of anesthetic (7.2% Chloralhydrate 10ml/kg. i.p.). Upon respiratory arrest, the animals were perfused transcardially with 200ml phosphate-buffered solution, followed by 200ml 4% paraformaldehyde in 0.1 m phosphate buffer solution. Brains were removed and postfixed overnight in a 4% paraformaldehyde and 30% sucrose solution (1:1), followed by immersion in 30% sucrose solution for at least 24 hours. The correct placement of the electrode was histologically verified in Nissl-stained sections.

### 2.8 Statistics

Statistical analyses were performed using SigmaStat 4.0 software (Systat Software Inc., San Jose, CA, USA). For behavioral testing, a two-way repeated measures analysis of variance (ANOVA) F-test with the factors group and time was used followed by a Bonferroni post hoc t-test for multiple comparisons. For the comparisons of the electrophysiological data between the deafened and the sham-deafened and non-operated control groups a Mann-Whitney U test was applied to test for significant differences. As the characteristics of the ISI histograms (mean, median, mode, and coefficient of variation) were not normally distributed in most cases, data were analysed with nonparametric multifactorial statistical methods. All tests were used two-sided; a p<0.05 was considered significant. GraphPad Prism 9 (GraphPAd Software, Inc 2020) was used to prepare the graphs.

## 3 Results

### 3.1 Surgery

All operated rats survived surgery without obvious behavioral disturbances and were reunited with their mother. One deafened animal kept losing weight in the following days and was therefore euthanized. Although in the following, all rats gained weight, the rats with hearing loss weighed about 10% less than the sham-deafened and naïve control groups, both during ad-lib feeding (until weight of 200g) and thereafter with controlled feeding of 16g/rat/day. Statistical analysis showed significant results for the factors group (F_2/647_=5.584 p=0.010) and week (F_23.647_=1099.373 p<0.001), with no interaction between factors (F_46.647_=0.882 p=0.693).

### 3.2 Hearing status

Intracochlear neomycin-injection at PND 14 lead to complete hearing loss (ABR threshold >120dB) in 8 rats. In four rats, complete hearing loss was observed on one side, whereas on the other side, only partial hearing loss was detected (two rats with 70 dB, one rat with 80 dB, and one rat with 90 dB). One sham rat was deaf on one side (120dB SPL), but had normal hearing on the other side, and was therefore excluded from further analysis. Thus, 12 deafened, 8 non-operated controls, and 7 sham-deafened rats were used for statistical evaluation. Final ABR at week 24 did not differ from initial measures. Sham-deafened and non-operated controls showed an ABR threshold of 40dB SPL. Statistical analysis with ANOVA showed a significant effect for the factor group (F_2,53_=146.103, p<0.001), while the factor time (F_1,53_=1.997, p=0.170) and interaction between factors (F_2,53_=0.447, p=0.645) was not significant. Post-hoc testing between groups showed enhanced ABR thresholds after neomycin injection compared to non-operated and sham-deafened controls (p<0.001; Figure 1B).

### 3.3 Behavior

#### Open field

In the open field, deafened rats covered more distance with increased velocity than non-operated or sham-deafened controls from postoperative week 8 on. For both measures, statistical analysis showed a significant interaction between the factors group and week (distance: F_10,161_=2.506, p=0.009; velocity: F_10,161_=2.635, p=0.006), with post-hoc testing showing significantly higher values in deafened rats from week 8 on, except for week 16 for the distance (all p<0.05; Figure 2A,B).

**Figure 2:**
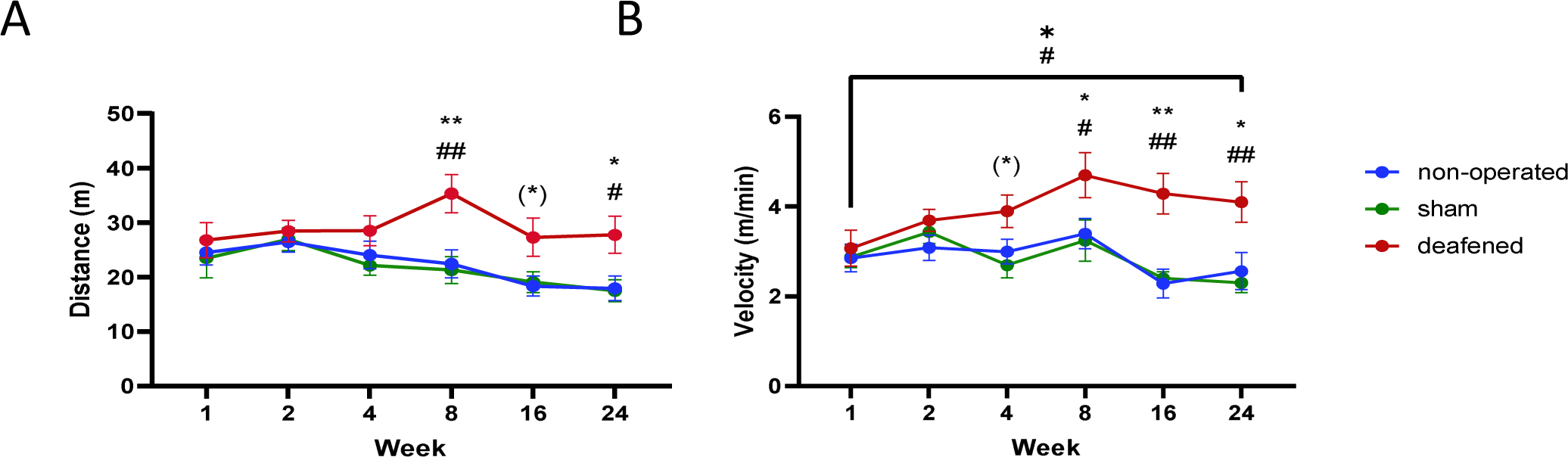
Open field. Results of testing for motor impairment in the open field during the experimental period. Distance travelled in the open field **(A)** and average velocity in the open field **(B)** with data shown as means ± SEM. Differences between deafened rats and non-operated controls are shown as asterisks (*) and between deafened and sham as hashtag (#) (*p < 0.05, **p < 0.01 after ANOVA); characters in brackets indicate significant intergroup-differences after trend for significance in ANOVA ((*) p < 0.1).

#### Social Interaction

The total duration of social interactions of deafened rats increased during adolescence, mainly due to increased play-fighting in this group. Statistical analysis of the total duration of all social interactions showed a significant effect for factor group (F_2,161_=10.025 p<0.001) and time (F_5,161_=6.650 p<0.001), but no interaction between factors (F_10,161_=0.712 p=0.712). Subsequent post hoc testing between groups showed increased overall social interaction in deafened rats as compared to sham-operated and non-operated controls (p=0.009), with no difference between the two control groups (p=1.0; Figure 3A). Analysis of play-fighting showed a significant effect for the factor group (F_2,161_=15.733 p<0.001), the factor week (F_5,161_=17.267 p<0.001) and the interaction between factors (F_10,161_=2.642 p=0.006). Post-hoc testing on the different weeks showed a significant increase in play fighting in deafened rats up to week 8 (all p<0.05), which was reduced by weeks 16 and 24, but still above that of sham-operated and non-operated control rats (Figure 3B). In contrast, statistical analysis of the duration of following behavior showed no significant effect for the factor group (F_2,161_=0.174 p=0.842) and the interaction between factors group and week (F_10,161_=1.065 p=0.395), but a significant effect for the factor week (F_5,161_=11.270 p<0.001), as the duration of following behavior decreased in all groups over the course of the study (Figure 3C).

**Figure 3:**
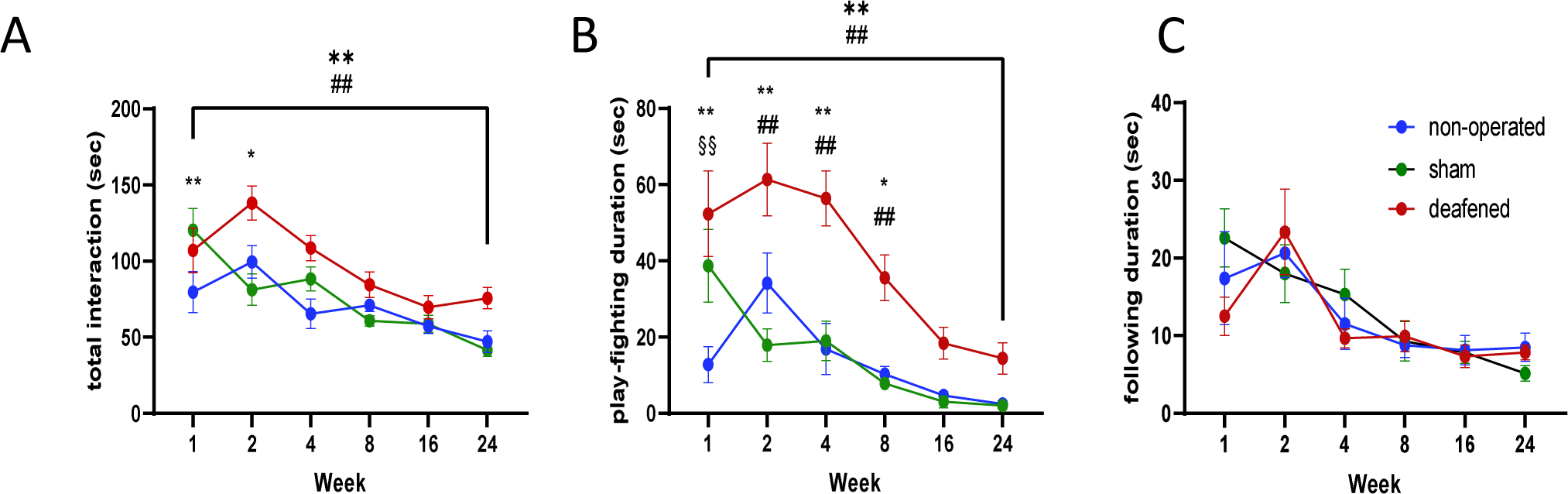
Social interaction. Total time of social interaction **(A)**, partial time of play-fighting **(B)**, partial time of following **(C)** each for the entire experimental period with data shown as means ± SEM. Differences between deafened rats and non-operated controls are shown as asterisks (*), between deafened and sham as hashtag (#) and differences between non-operated and sham as section sign (§) (*p < 0.05, **p < 0.01 after ANOVA).

#### Social preference

In the social preference paradigm, all animals preferred the filled (social partner) over the empty cage (object) with no differences between groups. Statistical analysis showed a significant effect for the factor duration of contact with empty/filled cages for all weeks (all F_1,53_>8.470, all p<0.008), whereas the factor group and the interaction between factors indicated no effect (all F_2,53_<1.290, all p>0.294; Supplementary Table 1).

#### Rotarod

Rotarod testing revealed no balance impairments in any of the experimental groups. Statistical analysis did not show any significant difference of the factors group (F_2,161_=1.762 p=0.193), time (F_5,161_=1.143 p=0.341) and interaction between factors (F_10,161_=0.640 p=0.777).

#### Plus maze

Rats tested in the plus maze did not show a preference for open or closed arms in the group comparison, but deafened rats showed an overall higher activity. The entries in open arms analyzed with a one-way analysis of variance showed no group difference (F_2,26_=1.448 p=0.255). The only significant result shown between deafened and both control groups was a strong increase in activity of the deafened rats by the total number of arms visited for factor group (F_2,26_=3.425 p 0.049) using a one-way ANOVA (Table 1).

**Table 1:** Plus maze. Total entries into closed and open arms and the resulting crossings of the central platform, split by deafened, sham, and non-operated rats. Differences between deafened rats and non-operated controls are shown as asterisks (*), between deafened and sham as hashtag (#) (*p < 0.05, after ANOVA); characters in brackets indicate significant intergroup-differences after trend for significance in ANOVA ((*) p < 0.1).

| Group | closed arm | open arm | central platform |
| --- | --- | --- | --- |
| **non-operated** | 17,38 ± 1,18 | 8,25 ± 1,70 | 25,63 ± 2,30 |
| **sham** | 12,86 ± 1,50 | 11,00 ± 0,98 | 23,86 ± 5,11 |
| **deafened** | 18,83 ± 1,32 # | 13,00 ± 1,40 | 31,83 ± 2,33 (#) |

#### 4-arm baited 8-arm maze

All groups learned the 4-arm baited 8-arm maze paradigm. During training, the number of total errors and the number of RME did not differ between groups, the number of WME was only initially increased during training in the deafened group. However, the 45° turns, as measure for next arm entries, were significantly reduced in the deafened group. Overall, there were no group differences in any of the parameters at the 16^th^ and 24^th^ week retests and the final reversal test.

As all rats learned the task during training, analysis with two-way RM ANOVA showed a significant effect for the factor “block” for all parameters, which is therefore not reported in the following. For training statistical analysis showed no group differences in total number of errors and the RME (total errors: F_2,26_=0.185 p=0.832; RME: F_2,404_=1.348 p=0.279; Figure 4A,B). Analysis of the WME showed a trend towards significance for the factor group (F_2,404_=2.876 p=0.076), with significantly more WME in the deafened group than in controls for block one and two (both p<0.016; Figure 4C). The deafened rats made less neighbor arm entries (45° turns) than control groups, as shown by a significant effect for the factor group (F_2,539_=4.390 p=0.024; Figure 4D).

**Figure 4:**
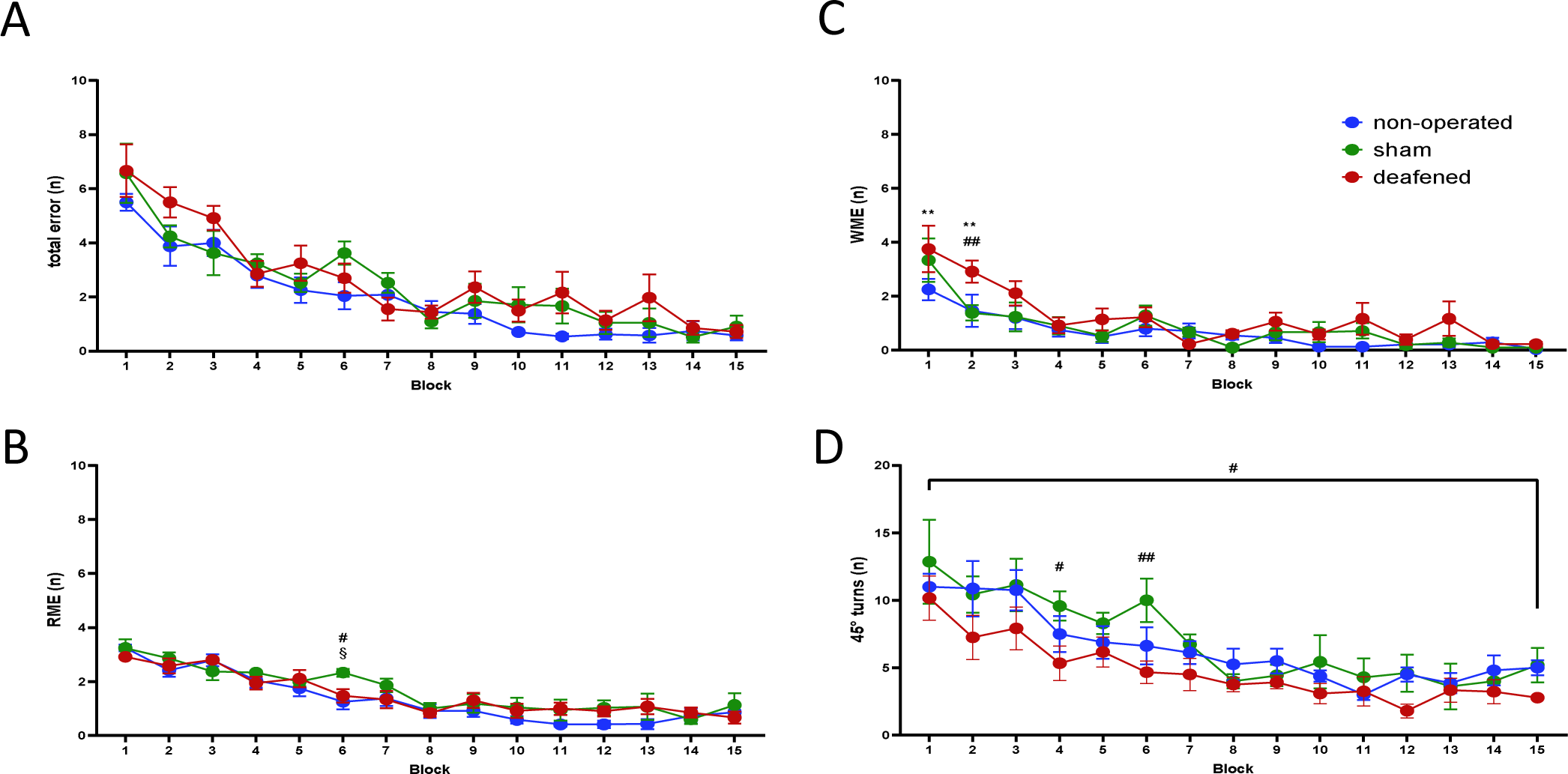
4-arm baited 8-arm maze. Total errors of rats initially learning the 4-arm baited 8-arm maze at week 8 after surgery for the different test blocks **(A)**, reference memory errors (RME) **(B)**, working memory errors (WME) **(C)** and 45° turns **(D)** with data shown as means ± SEM. Differences between deafened rats and non-operated controls are shown as asterisks (*), between deafened and sham as hashtag (#) and differences between non-operated and sham as section sign (§) (*p < 0.05, **p < 0.01 after ANOVA).

Overall, groups performance at weeks 16 and 24 and the final reverse test showed no differences between groups for total errors, RME, WME, and 45° turns (all p>0.1). Only at week 16, deafened rats made more RMEs than the other groups (F_2,107_=3.427 p<0.049; Supplementary Figure 1).

### 3.4 Electrophysiology

For electrophysiological analysis, spontaneous SU activity was recorded in the mPFC with a total number of n=658 (65.8 ± 9.5 per rat) for the deafened group, n=448 (64 ± 8.4 per rat) for the sham-operated controls, and n=466 (58.3 ± 8.2 per rat) for the non-operated controls. Analysis of the waveform durations allows to classify neurons as either NS for GABAergic as inhibitory neurons and BS for glutamatergic as excitatory neurons. A 10-second recording epoch is shown as an illustrative example of the extracellular spikes of a single neuron and their raster plots from the mPFC along with the action potential waveform (Figure 5). For neuron analysis we used recorded neurons in the mPFC and divided them into BS and NS for the non-operated controls (BS n = 378, NS n = 88; relation: 81.1/18.9), the sham-deafened group (BS n = 349, NS n= 99; 77.9/22.1), and the deafened group (BS n = 548, NS n = 110; relation: 83.3/16.7). The average frequency of NS (mean and SEM) recorded per individual rat in the non-operated group in the mPFC was 6.80 ± 0.49, for sham-operated rats 6.50 ± 0.45 and for deafened rats 4.91 ± 0.32. For the BS, the average recorded per individual rat for the non-operated rats was 6.25± 0.20, for the sham-deafened 6.59 ± 0.20, and for the deaf 5.62 ± 0.15.

**Figure 5:**
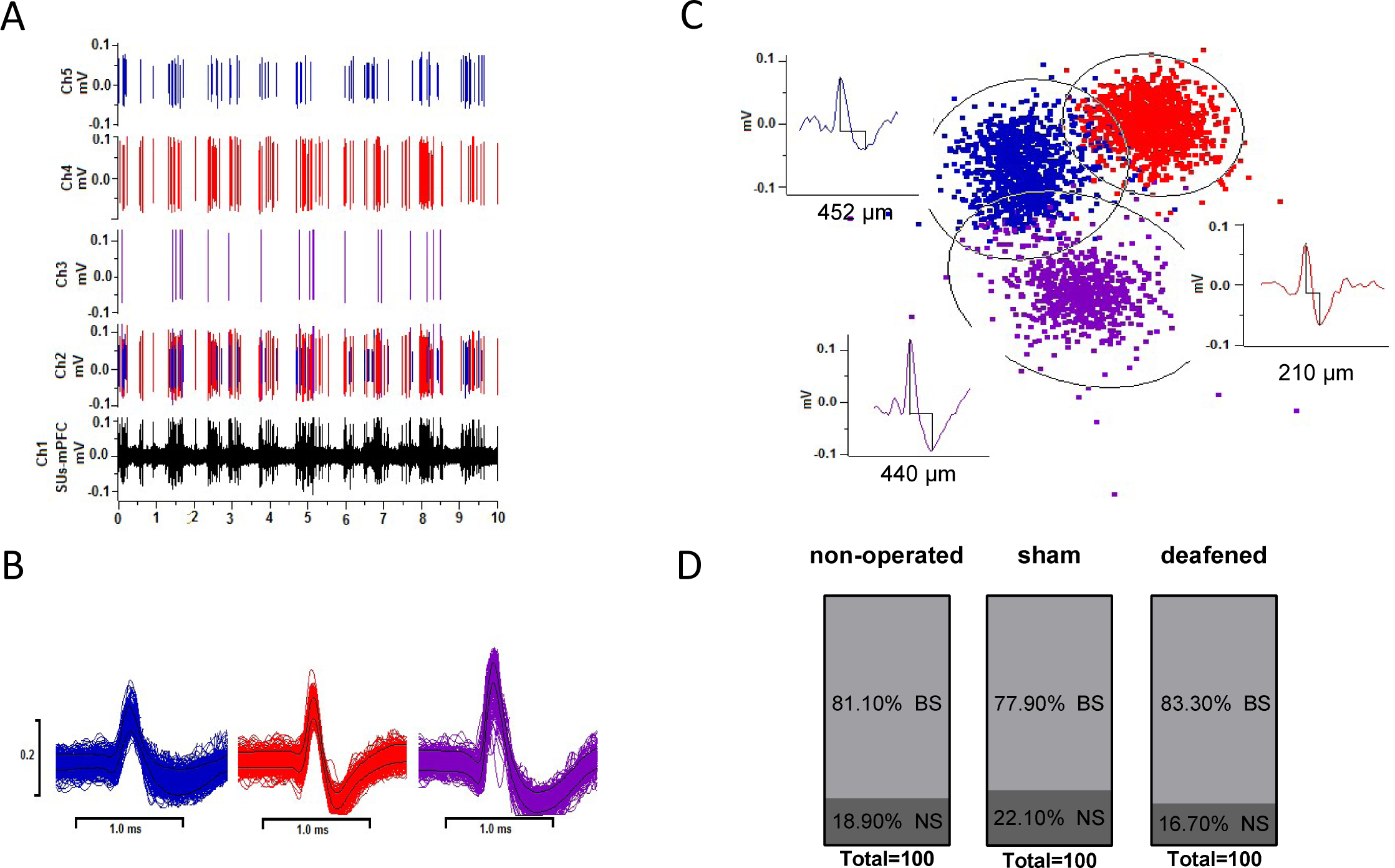
Neuronal spike sorting. Example of a 10-s recording epoch of the raw extracellular neuronal spikes from the mPFC (channel 1 in black). Channel 2 shows the raster plots with all different types of single neuronal units identified by spike sorting. Individual differentiated spikes from single units recorded in the mPFC are shown above (channels 3, 4, and 5) **(A)**. Corresponding associated action potential waveforms are shown in the panels below **(B)**. On the right side are examples of short duration spikes (peak to trough time of 0.12-0.36 ms), considered as narrow spike (NS) interneurons, and long duration neurons (time window >0.36 ms), considered as broad spike (BS) pyramidal neurons **(C)**. Furthermore, the percentage distribution of BS and NS in the respective groups is presented **(D)**.

The FR in the mPFC of deafened rats was lower than in the sham-operated and the non-operated control groups (both p<0.001; Figure 6A), also when classified into broad and narrow spike SU activity (p<0.049 and p<0.020, respectively; Figure 6B, C). The total dispersion index of deafened rats was increased compared to the sham-operated rats (p<0.001; Figure 6A), which did not reach the level of significance compared to the non-operated controls (p=0.078). When classified into broad and narrow spike SU activity, the dispersion index was only higher in the broad spike activity of deafened rats compared to sham-operated rats (p<0.001), but not to non-operated controls (p=0.764). No group differences were found for the analysis of NS (Figure 6B, C). Analysis using a one-way ANOVA showed significantly lower mean frequencies in NS and BS (both p<0.031) with no difference between the control groups (p>0.640).

**Figure 6:**
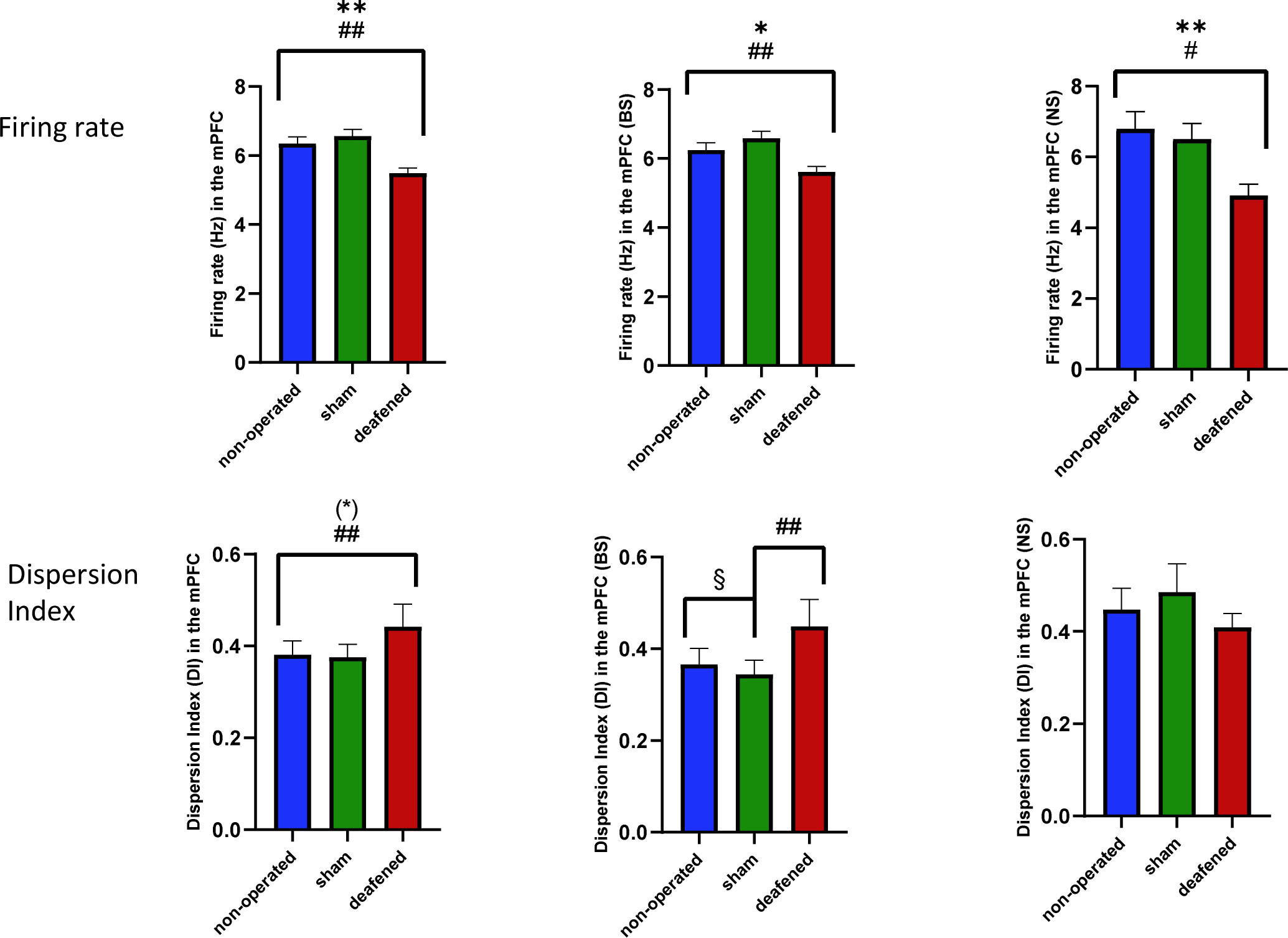
Single unit activity. Single unit activity for which both firing rate and disperson index were measured. Results are plotted as total **(A)** and then further differentiated into broad spikes for putative pyramidal neurons **(B)** and narrow spikes for GABAergic inhibitory neurons **(C)**. Results of deafened, sham and non-operated are shown as mean ± SEM. Differences between deafened rats and non-operated controls are shown as asterisks (*), between deafened and sham as hashtag (#) and differences between non-operated and sham as section sign (§) (*p < 0.05, **p < 0.01 after ANOVA); characters in brackets indicate significant intergroup-differences after trend for significance in ANOVA ((*) p < 0.1).

For analysis of the LFPs in the mPFC for the non-operated group n= 88, for the sham-deafened group n=76, and for the deafened group n=192 epochs were used. For the analysis of the relative power in SMCtx, n=146 samples were used for the deaf group, n=119 for the non-operated group, and n=72 samples for the sham-deafened group.

The one-way ANOVA analysis on the relative power of LFPs in the mPFC showed a significantly higher theta and alpha band activity compared to both control groups (non-operated: p<0.001 and sham-deafened: p<0.018). Beta and gamma band activity were lower in deafened rats compared to the non-operated control group (p <0.001). Further, beta band was also lower in sham-deafened as compared to the non-operated group (p<0.001; Figure 7A). Analysis of the relative power in SMCtx showed that only theta band activity was lower in deafened as compared to the sham-deafened and non-operated control groups (both p<0.031; Figure 7B).

**Figure 7:**
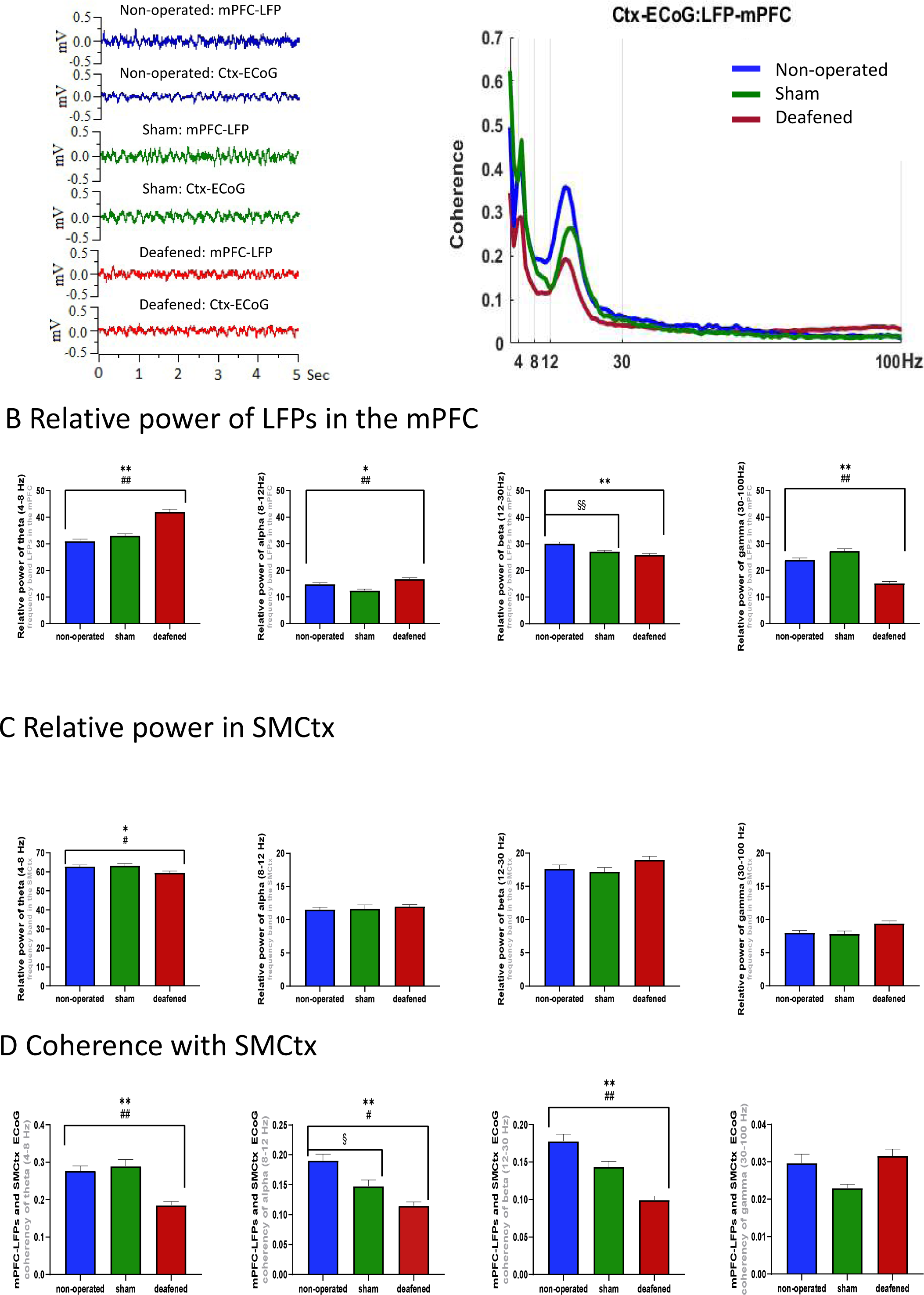
Local field potentials and coherence. Raw LFP signals as well as the average coherence spectrum within frequency range (4–100 Hz) across mPFC LFPs and sensorimotor cortex ECoG (A). The relative power of LFPs recorded from mPFC (B), the SMCtx (C) and the coherence between mPFC and SMCtx (D) of non-operated, sham and deafened rats are shown as mean ± SEM. **From left to right:** Recording of oscillatory activity of theta (4-8Hz), alpha (8-12Hz), beta (12-30Hz) and gamma (30-100Hz) frequency bands. Differences between deafened rats and non-operated controls are shown as asterisks (*), between deafened and sham as hashtag (#) and differences between non-operated and sham as section sign (§) (*p < 0.05, **p < 0.01 after ANOVA).

The coherence analysis with SMCtx-ECoG showed significantly reduced activity in deafened rats in theta (both p<0.001), alpha (both p<0.015) and beta (both p<0.001) frequencies compared to the two control groups, while gamma did not differ. Only alpha activity was reduced in sham-deafened as compared to the non-operated group (p=0.028; Figure 7C, D).

### 3.5 Histology

With the HE-color-series, the correct localization of the microelectrode used in electrophysiological leads could be accurately determined in the mPFC.

## 4 Discussion

Deaf and hard of hearing children are at higher risk of poor socio-emotional development and cognitive problems, which may persist in about half of the children after typical early intervention for hearing impairment, despite substantial improvements in language. It has been proposed that early auditory deprivation alters functional connectivity within the auditory system and also higher-order neurocognitive centres, resulting in risks of behavioral deficits, including executive function and concept formation (Kral et al., 2016). Less clear is whether the association between hearing loss and cognitive function is direct or indirect, e.g. through language competence (Hall et al., 2017). To gain insight into this question, we investigated the long-term behavioral effects of hearing loss induced in juvenile rats and the consequences for neural activity in the PFC in an experimental confined setting. Indeed, we found abnormal play in hearing impaired animals, that may be related to abnormal activity in prefrontal cortical neurons.

Juvenile rats normally engage in rough and tumble social play behavior, which increases until about 36 days of age (Thor & Holloway, 1984). Play has a key role in development of executive function (Yogman et al., 2018) and prefrontal cortex (Stark et al., 2023). Deficits in this behavior have been associated with neurodevelopmental disorders (Papilloud et al., 2018). The pronounced play-fighting in deafened rats, which persisted beyond the normal course of development, is therefore one of the most intriguing behavioral findings of the present study, especially since other aspects of social interaction did not differ between the groups and no obvious signs of enhanced aggression was seen. Taken together, these data suggest an association of hearing loss to play behavior through hyperactivity.

Deafened rats also showed a delayed onset of hyperactive behavior, as evidenced by an increased distance and speed in the open field from two months on after deafening. Another indication of hyperactivity is the increased total number of arms visited during the plus maze test. These observations compare well to the increased impulsivity and problems with executive control observed in late-implanted deaf children (Dye & Hauser, 2014; Yucel & Derim, 2008), who attributed more attention on visual periphery (Lomber et al., 2010; Neville & Lawson, 1987) and less to center, leading to deficits in sustained attention (Dye & Hauser, 2014; Yucel & Derim, 2008). It is also interesting to note that the deafened rats gained less weight, both during ad lib and during controlled feeding, than the control groups. Apparently, deafened rats consume more energy due to their hyperactivity. The combination of increased activity and play behavior seen after deafening has been used as an endophenotype for hyperactivity disorder in rodent models (Magnin & Maurs, 2017; Regan et al., 2022). An alternative explanation could be that these behaviors reflect an adaptive strategy to access information from the environment that is not available through hearing. Interestingly, hearing loss induced in adult rats had no effect on motor activity except for a transient increase in the distance walked in the open field one month after deafening (Johne et al., 2022). On the other hand, adult deafened rats were impaired on the Rotarod, a test traditionally used to assess balance deficits in rodents (Brooks et al., 2012; Hamm et al., 1994), while juvenile deafened rats did not differ from sham- and naïve controls. Apparently, due to the greater regenerative capacity of younger animals, some deficits can be easily compensated for, while others lead to compromised functional integrity of neuronal networks with behavioral, neurophysiological, and neuroanatomical consequences (Harich et al., 2008; Helgers et al., 2020).

In the 4-arm baited 8-arm radial maze test, optimal performance requires the rat to learn and to remember which arms are baited (reference memory component) and also which arms have been visited previously (working memory component). Typically, rats initially use an egocentric “procedural” strategy during training, i.e. they use their body position to guide their response by always turning right or left, resulting in enhanced 45° turns or next arm entries (Klein et al., 2004). Later on, this behavior is replaced by a more appropriate allocentric strategy, i.e., rats skip unbaited arms. This strategy requires the rats to create a central representation of external cues, a so-called “spatial map” of the environment. During training for the radial maze task in the present study, juvenile deafened rats made slightly more errors in the early learning phase, which is similar to that described after adult deafening (Johne et al., 2022). However, while the control groups readily used the next arm entry strategy, the deafened rats chose a more random search strategy, resulting in fewer entries into the next arm. This finding may indicate a reduced capacity for procedural learning, including a tendency to make suboptimal decisions as seen in patients with hyperactivity disorders (Sonuga-Barke & Fairchild, 2012).

Taken together, our behavioral findings replicate at least some aspects of the higher prevalence of hyperactivity, impulsivity and socioemotional problems described in children with hearing impairment (Altshuler, 1978; Eldik, 2005; Fellinger et al., 2009; González-Bautista et al., 2021; Theunissen et al., 2014; Van Eldik et al., 2004). Alterations in brain plasticity are a known factor in these behavioral disorders (Casey & Durston, 2006; Rapoport & Gogtay, 2008; Shaw et al., 2007). As the mPFC plays a cardinal role in higher cognitive abilities, socio-emotional and executive functions, including the formation of abstract categories and concepts (Teffer & Semendeferi, 2012), we focused here on the electrophysiological analysis of neural activity in this region.

In the neocortex, excitatory pyramidal neurons are the most abundant cell type, which can project across brain areas, while interneurons are predominantly inhibitory and their axons remain within a circumscribed cortical area, indicating their function as local processing units (Diester & Nieder, 2008). Electrophysiological recordings of SU activity and LFPs in the mPFC were performed in rats 6 months after hearing loss to identify possible long-term neural changes. In the present study and similar to adult deafened rats (Johne et al., 2022), the firing rate in the mPFC of juvenile deafened rats was reduced along with an increased irregular firing. Although this pattern did not differ between glutamatergic and GABAergic neurons, the ratio of these two types of neurons was somewhat altered with fewer inhibitory GABAergic neurons in juvenile deafened rats, which may lead to an imbalance of neuronal activity within the mPFC and inefficient output to other regions.

In addition to SU activity, oscillations and rhythmic activities of neural networks within and between brain regions are fundamental for complex perceptual and cognitive functions, including language and social communications (Murphy & Benítez-Burraco, 2017; Tendler & Wagner, 2015). In the present study, oscillatory theta and alpha band activity in the mPFC was enhanced, while beta and gamma band activity was reduced rats with hearing loss. In the SMCtx area, however, only theta band activity was reduced while other frequency bands were not affected. These findings indicate that hearing loss in rats may lead to alterations in the oscillatory activity and synchronization patterns specifically within the mPFC. Theta band activity is modulated by higher order cognitive processes such as working memory and often interacts with gamma band activity (Zielinski et al., 2020), often in synchrony with the hippocampus via its ventral subregion (O’Neill et al., 2013). Furthermore, theta band activity is altered in psychiatric disorders with concomitant cognitive impairment (Li et al., 2015).

Coherence analysis is commonly used to investigate functional connectivity and communication between brain regions. Our coherence results showed an overall decrease in the strength of theta, alpha, and beta synchronization, while the gamma frequency band remained unchanged. A higher coherence in the theta frequency band would indicate higher connectivity of information processing between two regions, suggesting efficient and better task performance (Duprez et al., 2020; Roux & Uhlhaas, 2014), whereas higher coherence in the alpha frequency band between the mPFC and sensorimotor cortex would be associated with inhibitory processes and attentional modulation (Foxe & Snyder, 2011; Klimesch, 2012; Wöstmann et al., 2017). The decrease in theta and alpha coherence in the deafened rat group may therefore indicate a decrease in working memory and inhibitory control, resulting in increased interference from irrelevant stimuli and a cognitive deterioration, as observed in the 4-arm baited 8-arm maze test during training. Beta frequency band reflects rhythmic processing (Gilley et al., 2016), that has been associated with sensory integration (Brovelli et al., 2004), working memory (Shahin et al., 2009; Zarahn et al., 2007), auditory template matching (Bidelman, 2017; Yellamsetty & Bidelman, 2018), and inhibitory processing (Kropotov et al., 2011) which is often impaired in neurodevelopmental disorders and movement disorders (Milner et al., 2018). Furthermore, abnormal gamma oscillations have been observed in various neurological and neurodevelopmental disorders (Vianney-Rodrigues et al., 2019). It has also been described in tinnitus in the auditory and somatosensory cortex (De Ridder et al., 2023). Nevertheless, gamma band activity was only reduced in the mPFC, but not in the SMCtx and not in coherence between mPFC and SMCtx.

Although altered neural activity in the mPFC likely reflect compensatory mechanisms, we hypothesise that it reduces its ability to recruit subsidiary brain regions and strategies for optimal behavioral outcomes. Adults with early-stage hearing loss showed additional recruitment of the frontal cortex, which is consistent with other studies showing increased activation of frontal cortices during degraded listening to situations in adults with normal hearing and adults with hearing loss. It has been proposed that these disturbances contribute to the deficits in cognitive and central auditory processing adults with cochlear implants (CI; (Kronenberger et al., 2013; Marschark et al., 2007). In parallel, frontal cortices are recruited in an effort to improve sensory perception through top-down modulatory control.

## 5 Conclusion

The present study links hearing loss to abnormal play behavior and abnormal prefrontal function. Furthermore, deficits in concept learning, as observed in rats, were similarly observed in deaf children (Kronenberger et al., 2014). Whether these are the direct consequence of hyperactivity and abnormal play behavior remains open. Despite marked improvements in spoken language outcomes long-term CI users are at increased risk for behavioral problems, such as hyperactivity and abnormal play behavior, together with deficits in complex cognitive skills (Kronenberger et al., 2013; Marschark et al., 2007). Here, we describe substantial evidence for impaired neuronal activity in the mPFC of juvenile deafened rats, which may contribute, at least in part, to the hyperactivity and neurocognitive deficits observed as long-term effects in absence of language. Although it is difficult to develop an animal model for all diagnostic dimensions of a behavioral change following hearing loss, our model reproduces distinct aspects and symptoms of this condition and may therefore be suitable for elucidating the mechanisms leading to this condition in more detail, and possible providing therapy.

## Data availability statement

The original contributions presented in this study are included in the article/Supplementary material, further inquiries can be directed to the corresponding author.

## Ethics statement

This animal study was reviewed and approved by Lower Saxony State Office for Consumer Protection and Food Safety.

## Author contributions

JJ, MJ, MA, JK, AK, and KS: study concept. JJ, MJ, MA, and KS: laboratory work and analyses. JJ, MJ and KS: scientific writing. All authors carefully revised the manuscript and approved the submitted version.

## Abbreviations

ABR: auditory brainstem response
AI: asymmetry index
ANOVA: analysis of variance
BS: broad spikes
DI: dispersion index
ECoGs: electrocorticograms
FR: Firing rate
ISI: inter-spike interval
LFP: local field potentials
mPFC: medial prefrontal cortex
NS: narrow Spikes
PND: postnatal day
RME: reference memory errors
SMCtx: sensorimotor cortex
SU: single unit
WME: working memory error

## Acknowledgments

We greatly appreciate the skilful technical assistance by Juergen Wittek and Monika van Iterson.

---

**Supplementary Figure 1:**
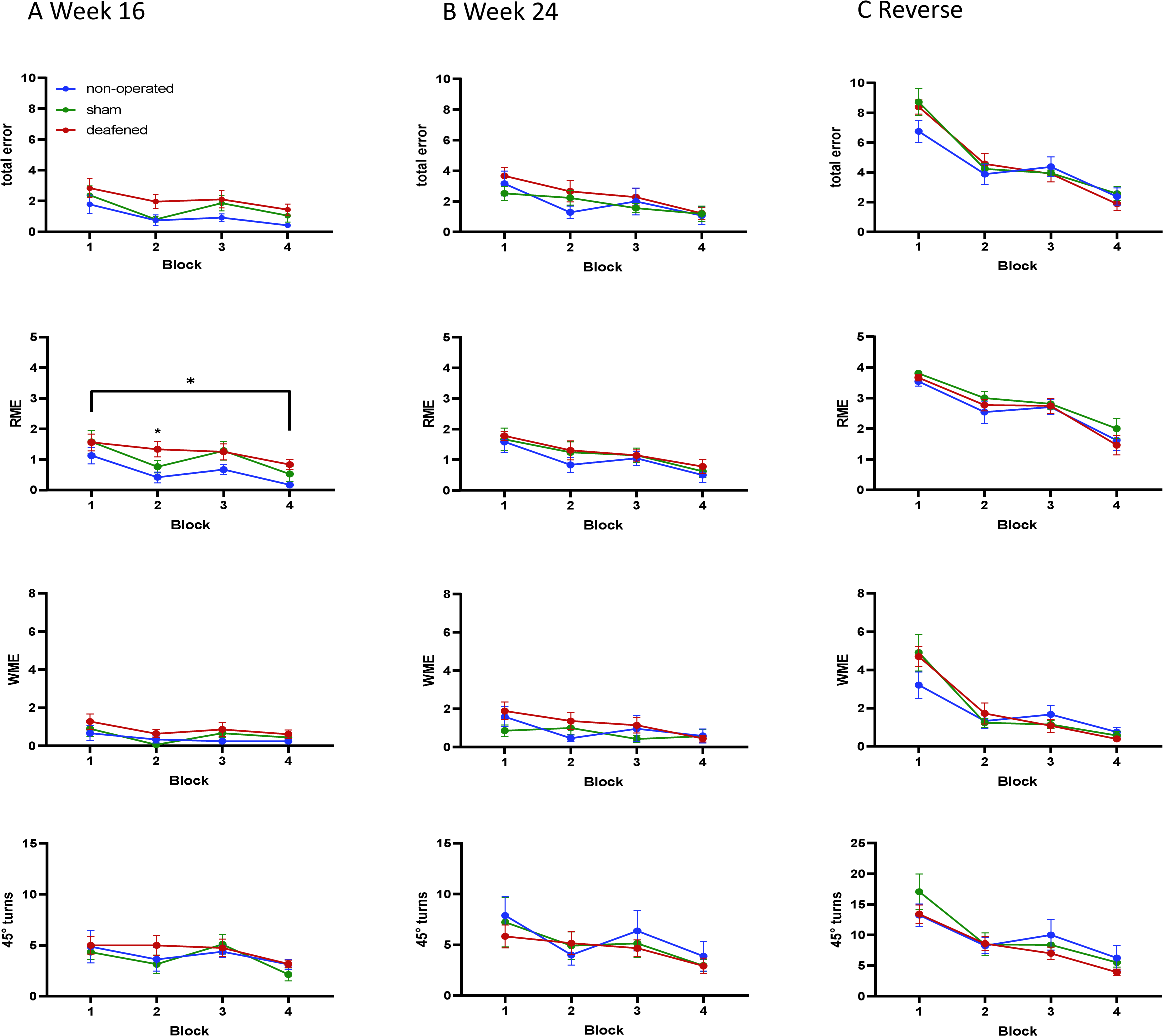
4-arm baited 8-arm. Total error of rats retested at weeks 16 **(A)** and 24 **(B)** and total error on the reversed test **(C)** in the 4-arm baited 8-arm maze shown as mean ± SEM. Included in descending order are the RME, WME, and 45-degree turns of the respective experimental section. Differences between deafened rats and non-operated controls are shown as asterisks (*p < 0.05).

**Supplementary Table 1:** Social preference. Total duration of interaction in the social preference paradigm with the empty containment compared to the containment filled with a social partner for all test weeks, broken down by experimental group as mean ± SEM. No significant differences were found after ANOVA.

|  | Week 8 |  | Week 16 |  | Week 24 |  |
| --- | --- | --- | --- | --- | --- | --- |
|  | empty | filled | empty | filled | empty | filled |
| non-operated | 111.5 ± 12.3 | 140.3 ± 12.3 | 89.5 ± 10.4 | 101.4 ± 10.4 | 72.8 ± 9.9 | 83.4 ± 9.9 |
| sham | 107.1 ± 13.2 | 145.4 ± 13.2 | 98.3 ± 11.1 | 118.1 ± 11.1 | 71.1 ± 10.6 | 97.6 ± 10.6 |
| deafened | 111.6 ± 10.1 | 142.2 ± 10.1 | 93.9 ± 8.5 | 109.6 ± 8.5 | 85.1 ± 8.1 | 92.3 ± 8.1 |

